# Comparative Genomic and Epigenetic Landscape of Histone H3.3 G34R and K27M-Mutated Pediatric High-Grade Glioma Suggests Divergent Precision Oncology Strategies

**DOI:** 10.64898/2026.09.14.751433

**Authors:** Zayan K. M. Islam, Lanlan Zhou, Wafik S. El-Deiry

**Affiliations:** Laboratory of Translational Oncology and Experimental Cancer Therapeutics, Warren Alpert Medical School, Brown University, Providence, RI, United States; Department of Pathology and Laboratory Medicine, Warren Alpert Medical School, Brown University, Providence, RI, United States; Legorreta Cancer Center at Brown University, Providence, RI, United States; Hematology/Oncology Division, Department of Medicine, Lifespan Health System and Brown University, Providence, RI, United States

**Keywords:** Glioma, DIPG, DMG, DHG, MGMT, ATRX, PDGFRA, SETD2, H3G34R, H3K27M

## Abstract

Histone *H3-G34R* and *H3-K27M* mutations define distinct diffuse high-grade glioma subtypes diffuse hemispheric glioma (DHG), midline (DMG), and pontine gliomas (DIPG). We characterized pathway-architectures using cBioPortal and pedcBioPortal for *H3-G34R* (n=112) and *H3-K27M* (n=727) mutated-gliomas, co-mutations, copy numbers (CNA), and variant data (n=112/727, n=86/578, n=51/313). We evaluated altered-genes as oncogenic by OncoKB, Gene-Ontology (GO) heatmaps and process-enrichment using clusterProfiler (Benjamini-Hochberg-adjusted p<0.05). *H3-G34R*-gliomas harbored near-universal *TP53* (94.1%), *ATRX* (81.4%) co-mutations and *PDGFRA* alteration (54.9% mutation; 17.1% amplification) within 4q12 amplicon that includes *KIT* and *KDR*. Pathway-enrichment converged on G1/S cell-cycle checkpoints, and epigenetic regulation, driven principally by *TP53*, *ATRX*, *PTEN*, *BCOR*, *TERT*, *FBXW7*, and *PPM1D*. *CDK6* amplification was rare (2.6%), despite *H3-G34R* association with CDK6-dependency including benefit from CDK4/6 inhibition. *H3-K27M*-gliomas displayed PI3K/AKT pathway activation (*PIK3CA*, 14.9%; *NF1*, 13.3%; *PDGFRA*, 10.8%, *PIK3R1*, 7.8%; *PTEN*, 5.7%) and developmental signatures through *NF1*, *PTEN*, and *SOX10* (4.95%) mutations. *TP53* (62.2%) and *ATRX* (21.2%) co-mutations were less frequent in *H3-K27M* versus *H3-G34R*, while *PPM1D* was *H3-K27M*-enriched (19.3%). *MGMT* promoter methylation occurred frequently in *H3-G34R* (65.2%, 15/23) versus *H3-K27M*-gliomas (5.4%, 4/74), despite genome-hypomethylation of *H3-G34R*. *H3-G34R*-gliomas, unlike *H3-K27M*, upregulated ganglionic *GSX2*, *DLX1*, *DLX2*, and *FOXG1*. *SOX10* was upregulated in *H3-K27M*-gliomas, versus *H3-G34R*. Lack of MGMT methylation in *H3-K27M*-gliomas correlates with temozolomide resistance; we previously reported imipridones ONC201/ONC206 reduce *MGMT*, *EZH1* and *EZH2* expression in *H3-K27M*-glioma cells. Divergent *PDGFRA*-mutations among *MGMT*-methylated versus unmethylated *H3-G34R*-gliomas warrants investigation of PDGFRA-signaling and epigenetic regulation. Our findings suggest divergent precision oncology therapeutic strategies to exploit unique vulnerabilities in *H3-K27M* and *H3-G34R*-gliomas.

## Introduction

Between 2017 and 2021, there were roughly 20,000 new cases of pediatric brain and central nervous system (CNS) cancers, leading to more than 2,000 deaths [1]. Brain and other CNS cancers are the leading causes of cancer-related death among pediatric patients, and pediatric high-grade gliomas (pHGGs), a group of highly aggressive tumors with dismal prognosis, are among the most notable subtypes [1–5]. pHGGs were officially established as a distinct, formal grouping for the first time in the 5th edition World Health Organization (WHO) classification of central nervous system tumors [6–9]. This reclassification reflected both the distinct underlying biology in pHGGs and their lack of response to existing treatment regimens for adult high-grade gliomas [6,7,9]. This new edition formally established histone *H3.3* mutation status as a defining diagnostic criterion for diffuse midline glioma (DMG) and diffuse hemispheric glioma (DHG), two of the four types of pHGGs [6–8]. DMGs are aggressive astrocytic tumors categorized as grade 4 malignancies [6–8,10]. In children, the most common manifestation of this tumor occurs in the pons and is referred to as diffuse intrinsic pontine glioma (DIPG) [1,6,11–13]. A lysine-to-methionine substitution at position 27 (K27M) in the H3.3 tail, most commonly in the *H3-3A* gene but occasionally in *H3-3B* or canonical *H3* genes such as *H3C2*, is the primary oncogenic driver of DMG [6,11,14,15]. Much like its counterpart, DHG is also an aggressive astrocytic grade 4 malignancy [6,7,9,14]. This tumor is predominantly located in the frontal and temporal lobes, and its primary oncogenic driver is the glycine-to-arginine substitution (G34R/V) at position 34 on the same H3.3 tail [6,7,9,14]. Although their oncogenic drivers are so similar, DMG and DHG co-occurrence of both mutations within a single tumor is incredibly uncommon [9,16].

Despite arising in anatomically and clinically distinct contexts, DMG and DHG share a similar epigenetic feature in that they both involve altered H3K27 trimethylation (H3K27me3) [9,12]. In DMG, K27M acts in trans, functioning as a dominant-negative inhibitor of the epigenetic regulatory complex PRC2’s catalytic subunit EZH2 and directly suppressing H3K27me3 deposition genome-wide [9,17,18]. In DHG, G34R/V instead acts in cis, impairing histone methyltransferase SETD2-mediated H3K36me3 [9,19,20]. Because the DNA methyltransferase DNMT3A, which reads H3K36me2/3 via its PWWP domain for chromatin recruitment, depends on this mark for chromatin recruitment, H3K36me2/3 loss both redistributes PRC2 activity and disrupts DNMT3A localization, driving a broader genome-wide DNA hypomethylation phenotype in DHG [9,20,21]. These convergent yet mechanistically distinct epigenetic programs have been proposed to arise from different developmental precursor cells-of-origin. Results from single-cell transcriptomic analysis suggest that DMG arises from an oligodendrocyte precursor cell (OPC) origin, and DHG a GABAergic interneuron progenitor origin within the ganglionic eminence [9,12]. Whether this divergence in proposed developmental origin is reflected in the somatic genomic architecture of each tumor – beyond the defining histone mutation itself – has not been systematically examined using a unified, directly comparable analytical pipeline across both subtypes. Along with this, *MGMT* promoter methylation, a determinant of response to temozolomide – which is standard first-line chemotherapy in DHG but of unproven benefit in DMG – has been reported as common in DHG but rare in DMG [9,19,22]. Specifically, we asked: (1) whether *H3-G34R* and *H3-K27M* gliomas exhibit distinct co-mutational signatures and pathway-level convergence points when analyzed using a matched, comparative genomic pipeline; (2) whether the well-documented divergence in *MGMT* promoter methylation between the two subtypes – which is paradoxical for *H3-G34R* given its genome-wide hypomethylation phenotype – shows any relationship to co-mutation patterns within each subtype; and (3) whether the somatic mutation landscapes of each subtype provide genomic evidence, independent of transcriptomic data, consistent with their proposed distinct cellular origins.

## Methods

### Data acquisition and cohort assembly

Mutation, copy-number alteration (CNA), and structural variant data (SV) for tumors harboring the *H3-G34R* and *H3-K27M* mutations were queried from cBioPortal for Cancer Genomics [23–25] and pedcBioPortal, an aggregated public repository of pediatric cancer genomic studies built on the cBioPortal platform. Sex-chromosome deletion was strongly concordant with each patient’s recorded sex (e.g., Y-linked gene loss was disproportionately called in patients recorded as female, X-linked in male), a common motif in genomics data [26,27]. As a result, genes located on sex chromosomes were excluded from CNA frequency analysis as suspected artifacts. Because no single contributing study provided a cohort of sufficient size for robust analysis, data were assembled across all available studies containing *H3-G34R* or *H3-K27M*-mutant samples on both cBioPortal and pedcBioPortal as of the query date (data pulled June-August 2026). The *H3-G34R* cohort comprised 94-samples from pedcBioPortal and 18-samples from cBioPortal (n=112 total with mutation data available); the *H3-K27M* cohort comprised 671-samples from pedcBioPortal and 56-samples from cBioPortal (n=727 total with mutation data available). Because CNA and SV profiling was not performed, or not reported, for every sample with mutation data, the amount of data on these properties differs from the mutation-data: CNA-profiled n=86 (G34R) and n=578 (K27M); SV-profiled n=51 (G34R) and n=313 (K27M). These assay-specific denominators are used throughout to compute frequencies and are reported alongside each figure and statistic. There were originally 729 *H3-K27M* glioma samples across both databases, but two additional sources of contamination were identified and excluded: one *H3-K27M*-annotated record corresponding to the cell line DIPG007 (study ccle_broad_2019, a cancer cell line dataset rather than patient tumor data), and one *H3-K27M*-annotated patient subsequently determined to have neuroblastoma rather than glioma, sourced from a non-glioma-specific pooled study (x01_fy16_nbl_maris). Both were excluded from all analyses in which they appeared.

### Pooling of fragmented sub-cohort data

Because the underlying data used for analysis was pooled across many contributing studies with different sample sizes, overall cohort frequencies were computed by summing the raw number of altered samples for each gene across all contributing sub-cohorts and dividing by the summed total number of profiled samples for the corresponding assay, rather than by averaging study-level percentages. This approach avoids the well-known bias of unweighted percentage-averaging, in which small studies are given disproportionate influence relative to their sample size. For patient-level clinical variables (e.g., age at diagnosis, overall survival), the same patient occasionally appeared under multiple study-version identifiers within the pooled pediatric portal data. Redundant records were deduplicated prior to analysis, ensuring that these data only considered each patient once.

### Tumor location analysis

Tumor location was extracted from two clinical attribute fields with differing nomenclature across contributing studies (TUMOR_TISSUE_SITE and Tumor Tissue Site), which were harmonized into a single location variable. Laterality (left/right) was not distinguished. Samples with a single reported anatomic site were classified into the corresponding hemispheric/cortical or midline/deep category; samples with two or more distinct named sites were classified separately as multifocal rather than counted toward each individual site category. To additionally capture diffuse site involvement obscured by multifocal classification, a secondary, non-mutually exclusive tally was computed reporting the number of samples with any involvement of each named site, irrespective of whether that site was the sole location or one component of a multifocal presentation

### Gene panel filtering

Recurrently altered genes identified in each cohort were filtered to those annotated as oncogenic cancer genes in the OncoKB precision oncology knowledgebase [28,29], restricting downstream pathway analysis to genes with established or likely relevance to tumorigenesis and reducing the influence of passenger alterations and artifacts common in large aggregated panels. For panels displaying recurrently altered genes, a default inclusion threshold of ≥5% alteration frequency was applied. An exception was made for the *H-3K27M* mutated-gene panel (**Figure 2B**), where a ≥4% threshold was used to retain *SOX10*, *FGFR1*, *ACVR1*, and *NOTCH1*. Although each of these genes were narrowly below the standard cutoff, they were included due to their established relevance to *H3-K27M* glioma biology. For the cross-cohort gene comparison (**Figure 3A**), the ten most frequently mutated genes from each cohort were selected and combined, such that a gene appears in the panel if it ranks among the top ten in either cohort.

**Table 1.** Cohort composition, demographics, tumor location, and assay-specific sample availability for *H3-G34R* and *H3-K27M* glioma cohorts in cBioPortal and pedcBioPortal. Values are n (%) unless otherwise noted. Some sample and profiling denominators differ from unique patient counts due to multiple sequenced specimens profiled for a subset of patients; alterations shared across a patient’s multiple samples may be overrepresented relative to a strictly one-sample-per-patient analysis. Age, sex, and tumor location data reflect patient-level counts, with no repeated contributions from multiple samples of the same patient; all other variables include repeat samples where applicable. For tumor location, 1° denotes classification to a single reported anatomic site (mutually exclusive across categories; sums with "Spans multiple sites" and "Unspecified/not reported" to the cohort location total); 2° denotes a non-mutually exclusive tally of any reported involvement of that site, including as one component of a multi-site entry, and therefore does not sum to the cohort total. "Other/not specified" indicates a specific but uncommon reported site not separately tabulated (e.g., basal ganglia, ventricles); this is distinct from "Unspecified/not reported," which reflects absent or non-informative location data and cannot be attributed to either the hemispheric/cortical or midline/deep cluster. All profiled samples carried a WHO grade 4 designations with one exception, a sample with an initial biopsy grade of 3, most likely reflecting limited-tissue sampling prior to definitive molecular classification.

|  | **H3G34R, (94 patients/ 112 samples)** | **H3K27M, (545 patients/ 727 samples)** |
| --- | --- | --- |
| Mutation | 112 | 727 |
| Copy-number (CNA) | 86 | 578 |
| Structural variant (SV) | 51 | 313 |
| Median age (years) | 15.0 | 9.0 |
| Age range | 7-58 | 1-47 |
| Male | 47 | 237 |
| Female | 31 | 268 |
| MGMT methylation + (%) | 15 (65.2%) | 4 (5.4%) |
| MGMT methylation - (%) | 8 (34.8%) | 70 (94.6%) |
| **Hemispheric/cortical** |  |  |
| Frontal (1°/ 2°) | 16 (26.7%) / 30 (50.0%) | 7 (1.5%) / 34 (7.3%) |
| Temporal (1°/ 2°) | 2 (3.3%) / 12 (20.0%) | 10 (2.2%) / 25 (5.4%) |
| Parietal (1°/ 2°) | 15 (25.0%) / 24 (40.0%) | 0 (0.0%) / 17 (3.7%) |
| Occipital (1°/ 2°) | 2 (3.3%) / 4 (6.7%) | 0 (0.0%) / 12 (2.6%) |
| Other/not specified | 0 (0.0%) | 0 (0.0%) |
| **Midline/deep** |  |  |
| Pons/brainstem (1°/ 2°) | 0 / 1 (1.7%) | 197 (42.5%) / 287 (62.0%) |
| Thalamus (1°/ 2°) | 0 / 2 (3.3%) | 66 (14.3%) / 150 (32.4%) |
| Spinal cord (1°/ 2°) | 0 / 1(1.7%) | 17 (3.7%) / 38 (8.2%) |
| Cerebellum (1°/ 2°) | 0 / 0 | 23 (5.0%) / 98 (21.2%) |
| Other/not specified | 0 / 3 (5.0%) | 9 (1.9%) / 66 (14.3%) |
| Spans multiple sites | 19 (31.7%) | 126 (27.2%) |
| Unspecified/not reported | 6 (10.0%) | 8 (1.7%) |

### Gene Ontology enrichment analysis

Gene Ontology (GO) Biological Process enrichment analysis [30,31] was performed separately on the OncoKB-filtered, recurrently altered gene sets for the *H3-G34R* and *H3-K27M* cohorts using clusterProfiler [32], querying the org.Hs.eg.db human genome annotation database [33] with Benjamini-Hochberg correction for multiple hypothesis testing (adjusted p<0.05). Semantically redundant GO-terms were collapsed via the simplify function, which is implemented using GOSemSim [34,35].

### Software

All data processing, statistical analysis, and figure generation were performed in R (v4.6.1) [36]. Data manipulation and wrangling were performed using the tidyverse collection of packages [37]. Figures were generated using ggplot2 [38], with GO enrichment dot plots produced using enrichplot [39] and composite multi-panel figures assembled using patchwork [40]. Anatomical maps of tumor location (**Figures 1F, 2F**) were generated using ggseg [41], using the Desikan-Killiany atlas for cortical/hemispheric regions and the ASEG atlas for subcortical/midline regions. Kaplan-Meier survival curves and log-rank tests (**Figure 3D**) were generated using the survival and survminer packages [42,43]. Wilcoxon rank-sum test annotations on comparative distributions (**Figure 3E, G, H**) were generated using ggpubr [44]. Molecular mechanism schematics (**Figures 1A, 2A, 5A**) were created using BioRender.com.

**Figure 1.**
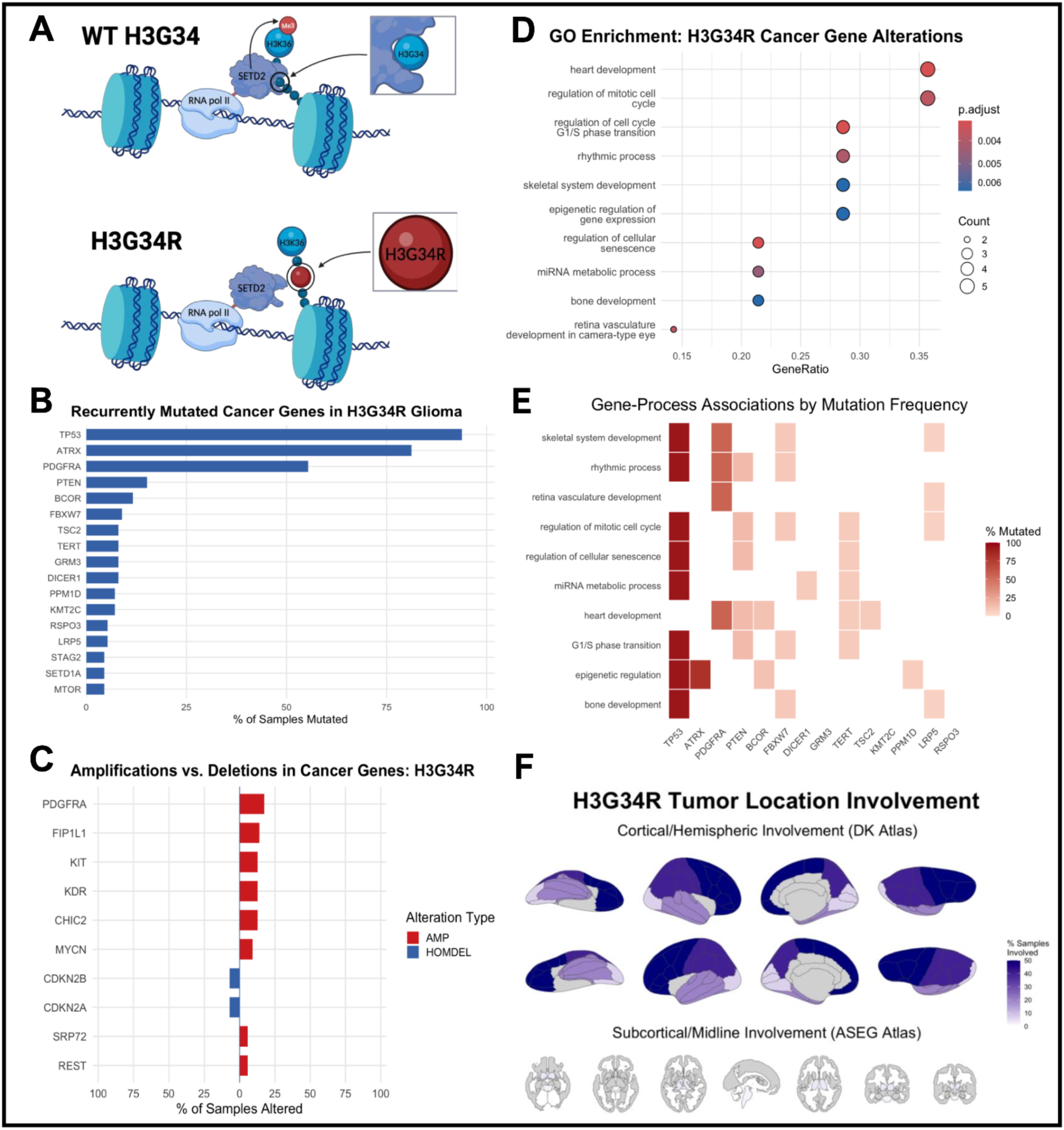
Genomic and epigenetic characterization of *H3-G34R*-glioma. **(A)** Schematic of SETD2-mediated H3K36 trimethylation under wild-type and G34R conditions. The G34R substitution sterically impairs SETD2 engagement with K36 in cis, reducing H3K36me3. This worsens DNMT3A recognition of H3K36 and results in altered genome-wide DNA methylation (Created with BioRender.com). **(B)** Frequency of OncoKB-filtered recurrently mutated cancer genes among mutation-profiled *H3-G34R* glioma samples. **(C)** Amplification (red) and homozygous deletion (blue) frequencies among recurrently altered cancer genes; genes on sex chromosomes were excluded due to ploidy-baseline artifacts in pooled copy-number calls. **(D)** Gene Ontology Biological Process enrichment among OncoKB-filtered mutated genes; dot size reflects gene count, color reflects Benjamini-Hochberg adjusted p-value. **(E)** Heatmap of enriched processes (rows) by contributing gene (columns); color reflects each gene’s overall cohort-wide mutation frequency, not a process-specific frequency. **(F)** Spatial distribution of tumor involvement by brain region, reflecting the non-mutually exclusive "any involvement" tally described in **Table 1**; subcortical/midline involvement was minimal (thalamus 3.3%, pons/brainstem 1.7%, spinal cord 1.7%, cerebellum 0%) and is not visually prominent at the color scale used for the cortical panel (up to 50%).

### MGMT promoter methylation and subgroup analyses

MGMT promoter methylation status was extracted from samples with available data for comparative subgroup analysis in both the *H3-G34R* and *H3-K27M* cohorts (G34R: 23 of 112 mutation-profiled samples; K27M: 74 of 727). Within each subtype, samples were stratified into *MGMT*-methylated and *MGMT*-unmethylated subgroups, and gene-level mutation and CNA frequencies were analyzed within each subgroup using the aforementioned pooling approach.

### Cell-of-origin gene panel

To evaluate genomic correlates of proposed cell-of-origin models, we examined mutation and CNA status, irrespective of OncoKB annotation, for a targeted panel of lineage-defining transcription factors: *SOX10* and *OLIG2* (oligodendrocyte precursor lineage, relevant to *H3-K27M*) and *GSX2*, *DLX1*, *DLX2*, and *FOXG1* (interneuron/ganglionic eminence lineage, relevant to *H3-G34R*), based on the developmental origin models proposed in prior single-cell transcriptomic studies [9,12]. We also investigated oligodendrocyte precursor lineage marker expression in *H3-G34R* tumor samples and interneuron/ganglionic eminence lineage marker expression in *H3-K27M* tumor samples to see if their comparative differentiation processes overlapped. Expression z-scores were queried from cBioPortal and pedcBioPortal, using RNA-seq (RSEM-normalized) data for *H3-G34R* and Agilent microarray data for *H3-K27M*; because these platforms differ, expression values are interpreted within, but not directly compared in magnitude across, each cohort. Where a sample contributed multiple expression records across overlapping contributing studies, values were averaged per sample prior to analysis; samples with a missing (NA) value for a given gene were excluded from analysis.

### Statistical considerations and limitations of approach

This study is a descriptive, hypothesis-generating comparative genomic analysis of publicly aggregated, cross-sectional cohort data and was not designed or powered as a formal hypothesis-testing study; consequently, we report frequencies with their exact numerators and denominators rather than p-values for most between-group comparisons, and we explicitly flag subgroup analyses with small sample sizes (n<20) as exploratory. An exception to this approach is the comparison of overall survival between the *H3-G34R* and *H3-K27M* cohorts (**Figure 3D**), for which a log-rank test was used given the substantially larger sample sizes available for this comparison relative to the other subgroup analyses in this study. Similarly, comparisons of age at diagnosis, fraction genome altered, and mutation count between the *H3-G34R* and *H3-K27M* cohorts (**Figure 3E, G, H**) used a Wilcoxon rank-sum test, given the larger sample sizes and continuous nature of these variables relative to the gene-level frequency comparisons that form the bulk of this study. No formal statistical test was used to distinguish candidate recurrent structural variant partners from background/noise-level findings; this interpretation instead relied on alteration frequency, the presence or absence of a clearly differentiated top candidate relative to other genes tested, and concordance with independently published structural variant findings in the relevant disease literature. The number of profiled samples exceeds the number of unique patients in both tumor cohorts (112 samples from 94 unique patients in the *H3-G34R* group; 727 samples from 545 unique patients in the *H3-K27M* group), reflecting that multiple specimens were sequence-profiled for some of patients. Because the pooled data were obtained as aggregate gene-level alteration frequencies rather than patient-linked sample records, restricting the analysis to a single sample per patient was not feasible. As a result, alterations shared across multiple samples from the same patient – most likely truncal or founder events – may be overrepresented relative to a strictly one-sample-per-patient analysis. This same limitation applies to the fraction genome altered and mutation count comparisons in **Figure 3G–H**, which were likewise computed at the sample level rather than deduplicated to one sample per patient. Public cBioPortal/pedcBioPortal aggregates draw on studies that used heterogeneous sequencing platforms, gene panels, and variant-calling pipelines, and clinical annotation (including *MGMT* methylation) was available for only a minority of samples.

## Results

### *H3-G34R* and *H3-K27M* glioma cohort overview in cBioPortal and pedcBioPortal

After pooling across contributing pedcBioPortal and cBioPortal studies, the *H3-G34R* cohort comprised 112 mutation-profiled samples (86 CNA-profiled, 51 SV-profiled) and the *H3-K27M* cohort comprised 727 mutation-profiled samples (578 CNA-profiled, 313 SV-profiled). Data-type availability, which varied both across and within cohorts, is summarized in Table 1. Median age at diagnosis was 15.0 years (range 7–58) in the *H3-G34R* cohort and 9.0 years (range 1–47) in the *H3-K27M* cohort. Both cohorts included a small number of adult-onset cases, consistent with reports that H3-mutant pediatric-type diffuse high-grade glioma, while predominantly associated with children and young adults, can rarely present in older adults [9,45]. Sex was reported for 78 *H3-G34R* patients (47 male, 31 female) and 505 *H3-K27M* patients (237 male, 268 female). *MGMT* promoter methylation status was available for 23 *H3-G34R* samples and 74 *H3-K27M* samples. Consistent with prior reports, *MGMT* methylation was common in *H3-G34R* (15 of 23, 65.2%) but rare in *H3-K27M* (4 of 74, 5.4%) [9,22].

Tumor location differed markedly between cohorts (Table 1, n=60 *H3-G34R*, n=463 *H3-K27M* samples with location data). Among *H3-G34R* glioma samples assigned to a single anatomic site, 35 of 60 (58.3%) were hemispheric/cortical, most commonly frontal (16, 26.7%) and parietal (15, 25.0%), while none were classified as purely midline/deep. The opposite pattern emerged in the *H3-K27M* cohort, where 312 of 463 (67.4%) single-site samples were midline/deep, dominated by the pons/brainstem (197, 42.5%) and thalamus (66, 14.3%), while only 17 of 463 (3.7%) were purely hemispheric. A substantial fraction of samples in both cohorts was annotated with more than one anatomic site (19 of 60, 31.7% in H3G34R; 126 of 463, 27.2% in *H3-K27M*). To capture site involvement obscured by this classification, we additionally computed a secondary, non-mutually exclusive tally in which each sample contributed to every site it involved, rather than being restricted to a single primary classification (**Table 1**, secondary columns). Although none of the *H3-K27M* glioma samples showed isolated parietal or occipital involvement in the primary analysis, this secondary analysis revealed 17 (3.7%) parietal-involving and 12 (2.6%) occipital-involving tumor sites in these samples. Pons/brainstem involvement rose to 287 of 463 samples (62.0%). Multi-site H3K27M entries involved more distinct structures on average than multi-site H3G34R entries (approximately 3.2 versus 2.2 sites per entry), consistent with the diffusely infiltrative growth pattern characteristic of DMG [4,46].

### *H3-G34R* Genomic and Epigenetic Characterization

The G34R substitution occurs near to lysine 36 (K36) on the H3.3 tail. Wild-type *H3-G34* encodes a glycine at residue 34, allowing the tail to fit into the catalytic domain on histone methyltransferase SETD2. However, the substitution to a bulky, positively charged arginine sterically hinders binding, and interferes with SETD2’s ability to engage and trimethylate K36 on the mutant histone specifically (**Figure 1A**). Because this defect is restricted in cis to the individual mutant H3.3 molecule rather than affecting SETD2’s catalytic activity broadly, the resulting loss of H3K36me3 is concentrated at genomic loci where mutant H3.3 is incorporated, rather than reflecting complete, genome-wide loss. However, the epigenetic landscape of *H3-G34R* tumors still presents widespread DNA hypomethylation, and the reduction in H3K36me3 impairs the recruitment of DNMT3A, a methyltransferase which reads H3K36me2/3 through its PWWP domain, resulting in genome-wide alterations in DNA methylation patterns [9,20,47].

Among OncoKB-filtered recurrently mutated cancer genes (**Figure 1B**), *TP53* (94.1%), *ATRX* (81.4%), and *PDGFRA* (54.9%) were the most frequently altered, consistent with prior reports [9,19,47]. A tail of additional recurrently mutated genes was observed at substantially lower frequency, including *PTEN, BCOR, FBXW7, TSC2, TERT, GRM3, DICER1, PPM1D, KMT2C, RSPO3, LRP5, STAG2, SETD1A*, and *MTOR*.

Copy-number analysis (**Figure 1C**) identified recurrent amplification of *PDGFRA* (17.1%) alongside co-amplification of *FIP1L1, KIT, KDR,* and *CHIC2*, all genes clustered within the same chromosome 4q12 locus known to co-amplify as a single amplicon rather than independently [6,48,49]. *MYCN* amplification was also observed. The only notable deletion events seen in this cohort were in the *CDKN2A/CDKN2B* locus at chromosome 9p21.3 (∼5%), a known tumor suppressor co-deletion thought to contribute to immunosuppression and poor prognosis across different cancers [50–52]. This finding supports previous findings on *CDKN2A/CDKN2B* alterations in *H3-G34R*-gliomas and provides biological reasoning for their reported sensitivity to CDK4/6 inhibitors [19,53]. Structural variant analysis of the *H3-G34R* glioma cohort did not identify a gene as a clear recurrent structural variant partner – the top candidates (*SETBP1, PCM1, GNAS, AKT3*) were low-frequency (∼6%) with no single gene clearly separating from the others in recurrence. Given the small cohort sample size for this analysis (n=51), a 6% frequency represents only 3/51 cases, and given the lack of connection to *H3-G34R*-glioma biology, these genes are likely a result of random rearrangements and not representative of a recurrent structural driver in this disease.

GO Biological Process enrichment based on the mutated genes list (**Figure 1D**) highlighted terms related to cell cycle regulation, cellular senescence, and epigenetic regulation of gene expression, plausibly driven by *TP53, PTEN, BCOR, FBXW7, TERT* and *ATRX*. **Figure 1E** maps each enriched process to its contributing genes, weighted by each gene’s overall mutation frequency, highlighting *TP53* and *ATRX* as the dominant drivers across most enriched terms. The GO function organizes terms in a hierarchical structure through a directed acyclic graph (DAG), where each term can have one or many parent (less-specific) terms and zero, one or many child (more-specific) terms accordingly [54]. Additional enriched terms reflected broader developmental processes, such as skeletal and bone development, linked to the Wnt pathway genes *RSPO3* and *LRP5*. Although these terms are not necessarily specific to glioma biology, their prevalence likely reflects the larger prominence of growth-related parent terms, and as such are interpreted accordingly in this context.

Consistent with the cohort-level tumor location findings (**Table 1**), spatial mapping of involvement (**Figure 1F**) confirmed *H3-G34R*-gliomas’ predominantly hemispheric/cortical distribution, most concentrated in frontal and parietal regions, with minimal subcortical/midline involvement.

### Characterization of *H3-K27M* Diffuse Midline Glioma

In wild-type cells, H3K27me3 occurs via deposition from EZH2, the catalytic subunit on Polycomb Repressor Complex 2, a group of genes involved in the epigenetic regulation of gene expression [55]. The *H3-K27M* mutation results in the lysine being swapped for methionine at position 27, and this mutation results in 22x higher affinity binding in EZH2’s catalytic pocket, resulting in non-productive, essentially inhibited protein [56]. This essentially leads to a quasi-sequestration of the PRC2 complex, reducing the genome-wide levels of H3K27me3 (**Figure 2A**). Unlike *H3-G34R*, which acts in cis by sterically impairing SETD2-mediated H3K36me3 on individual mutant nucleosomes, the epigenetic impact of *H3-K27M* at the protein level is in trans, meaning that it exerts its effect genome-wide.

**Figure 2.**
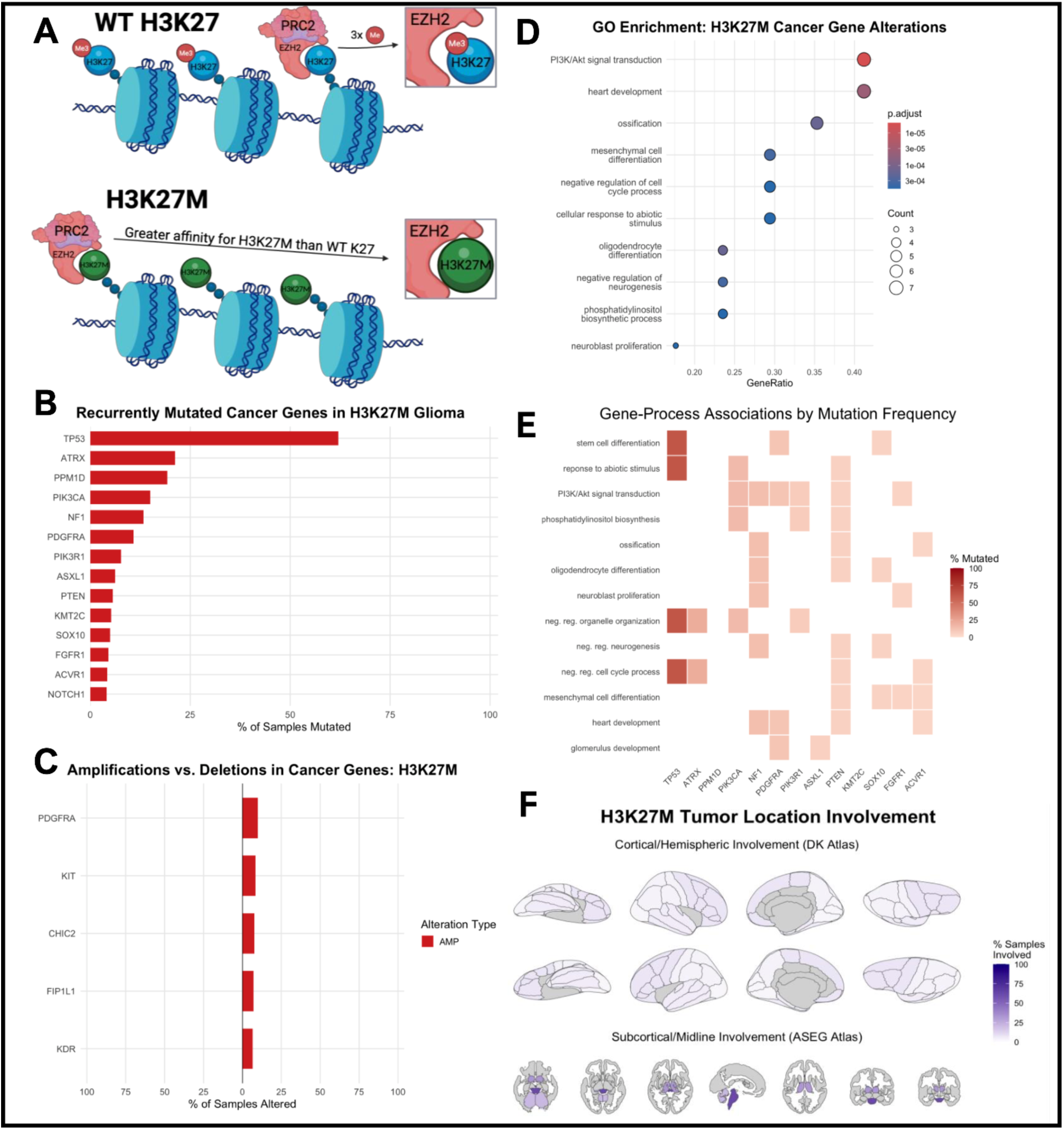
Molecular and genomic characterization of *H3-K27M* diffuse midline glioma. **(A)** Schematic of the H3-K27M mechanism. In wild-type H3K27 nucleosomes, PRC2 processively trimethylates H3-K27 via its catalytic EZH2 subunit. In H3-K27M-mutant nucleosomes, the mutant methionine residue binds EZH2 with a 22x greater affinity than wild-type H3-K27, trapping PRC2 and depleting the pool available to methylate wild-type H3-K27 elsewhere in the genome, resulting in global H3-K27 hypomethylation (Created with BioRender.com). **(B)** Recurrently mutated cancer genes, shown as percentage of profiled samples harboring a mutation (727 samples). Genes mutated in ≥4% of samples are shown. **(C)** Recurrent copy-number amplifications among cancer genes, shown as percentage of profiled samples (578 samples). **(D)** Gene Ontology Biological Process enrichment among recurrently mutated cancer genes. Dot size reflects gene count per term; color reflects Benjamini-Hochberg-adjusted p-value. **(E)** Heatmap of gene-process associations, colored by per-gene mutation frequency, showing which recurrently mutated genes contributed to each enriched GO term in **(D)**. **(F)** Spatial distribution of tumor involvement by brain region, reflecting the non-mutually exclusive "any involvement" tally described in **Table 1**; cortical/hemispheric involvement was minimal (frontal 7.3%, temporal 5.4%, parietal 3.7%, occipital 2.6%).

A group of 17-genes were recurrently mutated in ≥4% of profiled glioma samples (**Figure 2B**). *TP53* was the most frequently altered gene (62.2%), followed by *ATRX* (21.2%), *PPM1D* (19.3%), *PIK3CA* (14.9%), *NF1* (13.3%), and *PDGFRA* (10.8%). *PIK3R1, ASXL1, PTEN, KMT2C, SOX10, FGFR1, ACVR1,* and *NOTCH1* were each mutated in a smaller fraction of samples (∼4-10%). Genes were included at a 4% frequency threshold rather than a round 5% cutoff to facilitate the inclusion and discussion of certain biologically established *H3-K27M* glioma-associated drivers, such as *SOX10* (4.95%) and *ACVR1* (4.1%).

Copy-number analysis identified recurrent amplification of a gene cluster at the chromosome 4q12 locus – *PDGFRA, KIT, CHIC2, FIP1L1*, and *KDR* – each amplified in approximately 5-10% of profiled samples (**Figure 2C**). This finding is consistent with the shared co-amplicon architecture also observed in the *H3-G34R*-glioma cohort (**Figure 2C**), found in that exact group of genes on the same chromosome 4q12 amplicon.

The 17 recurrently mutated cancer genes were used as input for Gene Ontology (GO) Biological Process enrichment analysis, which identified significant enrichment (p.adjust < 0.05) for terms including PI3K/Akt signal transduction, oligodendrocyte differentiation, and negative regulation of neurogenesis, among others (**Figure 2D**). The enrichment of PI3K/Akt signaling and oligodendrocyte differentiation is notable, given the proposed oligodendrocyte precursor cell (OPC)-like origin of *H3-K27M*-altered diffuse midline glioma, and the important role of PI3K/Akt signaling pathways in the differentiation of OPC-like cells [57]. To identify which genes drove each enriched process, gene-process associations were visualized as a heatmap of per-gene mutation frequency (**Figure 2E**). *TP53* and *ATRX* contributed most heavily to negative regulation of cell-cycle process and negative regulation of organelle organization, while *PIK3CA, PIK3R1, NF1, PDGFRA, PTEN,* and *FGFR1* all converged on PI3K/Akt signal transduction, with *PIK3CA, PIK3R1*, and *PTEN* also being involved in the phosphatidylinositol biosynthetic process. Tumor location data showed strong concentration within midline/deep structures, particularly the pons/brainstem (**Figure 2F**), consistent with the classical DIPG presentation long associated with *H3-K27M*-altered gliomas.

### Comparative genomic and clinical analysis of *H3-G34R* and *H3-K27M* glioma cohorts

Recurrently mutated cancer genes were compared across both *H3-G34R* and *H3-K27M* glioma cohorts (**Figure 3A**). *TP53* and *ATRX* were mutated substantially more frequently in *H3-G34R* (94.1% and 81.4%, respectively) than in *H3-K27M* (62.2% and 21.2%), consistent with the established *TP53/ATRX/G34R* molecular signature of hemispheric high-grade glioma [9,19]. *PDGFRA* was likewise mutated more frequently in *H3-G34R* (54.9%) than *H3-K27M* (10.8%). Conversely, *PIK3CA, NF1*, and *PPM1D* were each mutated more frequently in the *H3-K27M* glioma cohort. CNA analysis identified a shared the chromosome 4q12 co-amplicon – *PDGFRA, KIT, CHIC2, FIP1L1,* and *KDR* – recurrently amplified in both cohorts, at modestly higher frequency in *H3-G34R* gliomas across all five genes (**Figure 3B**).

**Figure 3.**
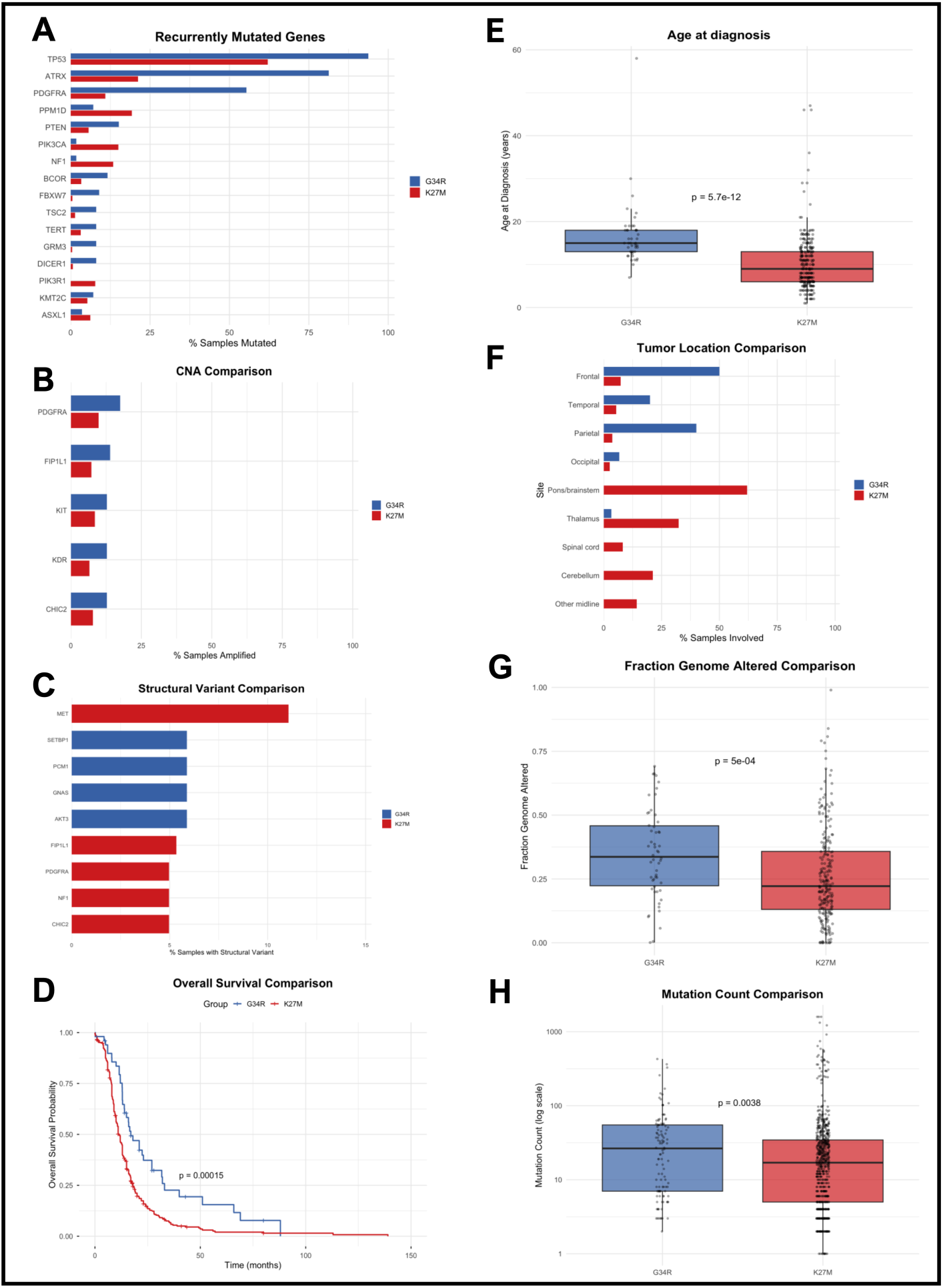
Comparative genomic and clinical analysis of *H3-G34R* and *H3-K27M* glioma cohorts. **(A)** Recurrently mutated cancer genes, restricted to the top 10 most frequently mutated genes in either cohort. **(B)** Recurrent copy-number amplifications shared between cohorts at the chromosome 4q12 locus. **(C)** Structural variant comparison. **(D)** Kaplan-Meier overall survival curves by cohort; log-rank test. **(E)** Age at diagnosis by cohort (*H3-G34R* n=45; *H3-K27M* n=260); Wilcoxon rank-sum test. **(F)** Non-mutually exclusive tumor location involvement by cohort, as percentage of samples with any reported involvement of each region. Percentages do not sum to 100% for either cohort; for *H3-G34R*, this reflects the combined effect of samples with unreported location (10%, contributing 0 to the tally) and samples with multiple reported sites (31.7%, contributing to more than one row), yielding a net total below 100% in this cohort. **(G)** Fraction genome altered by cohort (sample-level); Wilcoxon rank-sum test. **(H)** Mutation count by cohort (sample-level, log scale); Wilcoxon rank-sum test.

Structural variant analysis revealed that *H3-K27M*-gliomas showed recurrent rearrangements in *MET* (11.1%) and in *PDGFRA, FIP1L1, NF1,* and *CHIC2* (∼5%) (**Figure 3C**). The top *H3-G34R* glioma structural variant genes were *SETBP1, PCM1, GNAS,* and *AKT3*. Each variant was present roughly ∼6% of the time but is interpreted as having limited biological significance. *PDGFRA* was recurrently altered by three distinct mechanisms across the two cohorts: mutation (54.9%) and amplification (17.1%) in *H3-G34R*, and structural rearrangement (5.0%) in *H3-K27M*. These represent convergent but mechanistically distinct routes to *PDGFRA* dysregulation in *H3.3*-altered high-grade glioma.

Overall survival was significantly longer in the *H3-G34R* glioma cohort than the *H3-K27M* cohort (log-rank test, p=0.00015; **Figure 3D**), consistent with the comparatively more aggressive clinical course long associated with *H3-K27M*-altered diffuse midline glioma [3,19]. Patients with *H3-G34R*-mutant tumors were significantly older at diagnosis than those with *H3-K27M* (median 15.0 years, range 7–58, n=45 vs. median 9.0 years, range 1–47, n=260; Wilcoxon rank-sum test, p=5.7×10^-12^; **Figure 3E**).

Tumor location differed markedly between cohorts (**Figure 3F**): *H3-G34R* gliomas were predominantly cortical/hemispheric (frontal 50%, parietal 40%, temporal 20%) with minimal midline involvement (thalamus 3.3%, pons/brainstem 1.7%, spinal cord 1.7%, cerebellum 0%), while *H3-K27M* gliomas were overwhelmingly midline, dominated by the pons/brainstem (62.0%), thalamus (32.4%), and cerebellum (21.2%), with comparatively minimal cortical involvement (frontal 7.3%, temporal 5.4%, parietal 3.7%, occipital 2.6%). Non-mutually exclusive involvement percentages do not sum to 100% for either cohort; for *H3-G34R* specifically, this reflects the combined effect of samples with unreported location (10%, contributing 0 to the tally) and samples with multiple reported sites (31.7%, contributing to more than one row), which in this cohort yield a net total below 100%.

Fraction genome altered was significantly higher in *H3-G34R* than *H3-K27M* gliomas (median 0.33, IQR 0.23–0.45 vs. median 0.22, IQR 0.14–0.35; Wilcoxon rank-sum test, p=5.0×10^-4^; **Figure 3G**). Mutation count was likewise higher in *H3-G34R* than *H3-K27M* gliomas (Wilcoxon rank-sum test, p=0.0038; **Figure 3H**), though this comparison should be interpreted cautiously given heterogeneity in sequencing platform and panel depth across the pooled studies contributing to each cohort, which may partly drive raw mutation count differences independent of underlying tumor biology.

### Comparison of *MGMT*-methylated and *MGMT*-unmethylated subgroups among *H3-K27M* and *H3-G34R* gliomas

Despite there being a limited number of samples with *MGMT*-promoter methylation data, the results between cohorts were markedly different. Although only 23 of the 112 *H3-G34R* glioma samples had *MGMT*-data, 15 of the 23 (65.2%) *MGMT*-tested samples were methylated, compared to only 4 of 74 (5.4%) *MGMT*-tested *H3-K27M* glioma samples (**Figure 4A**). Given the small number of *MGMT*-methylated *H3-K27M* samples, group-level comparisons stratified by *MGMT* status were restricted to the *H3-G34R* cohort. Individual case data for the four *MGMT*-methylated *H3-K27M* samples are provided in **Supplementary Table 1 and 2**. Due to the small size of all cohorts with *MGMT* data, this analysis is strictly exploratory and is not meant to be interpreted as a definitive comparison of molecular or clinical correlates between *MGMT* subgroups.

**Figure 4.**
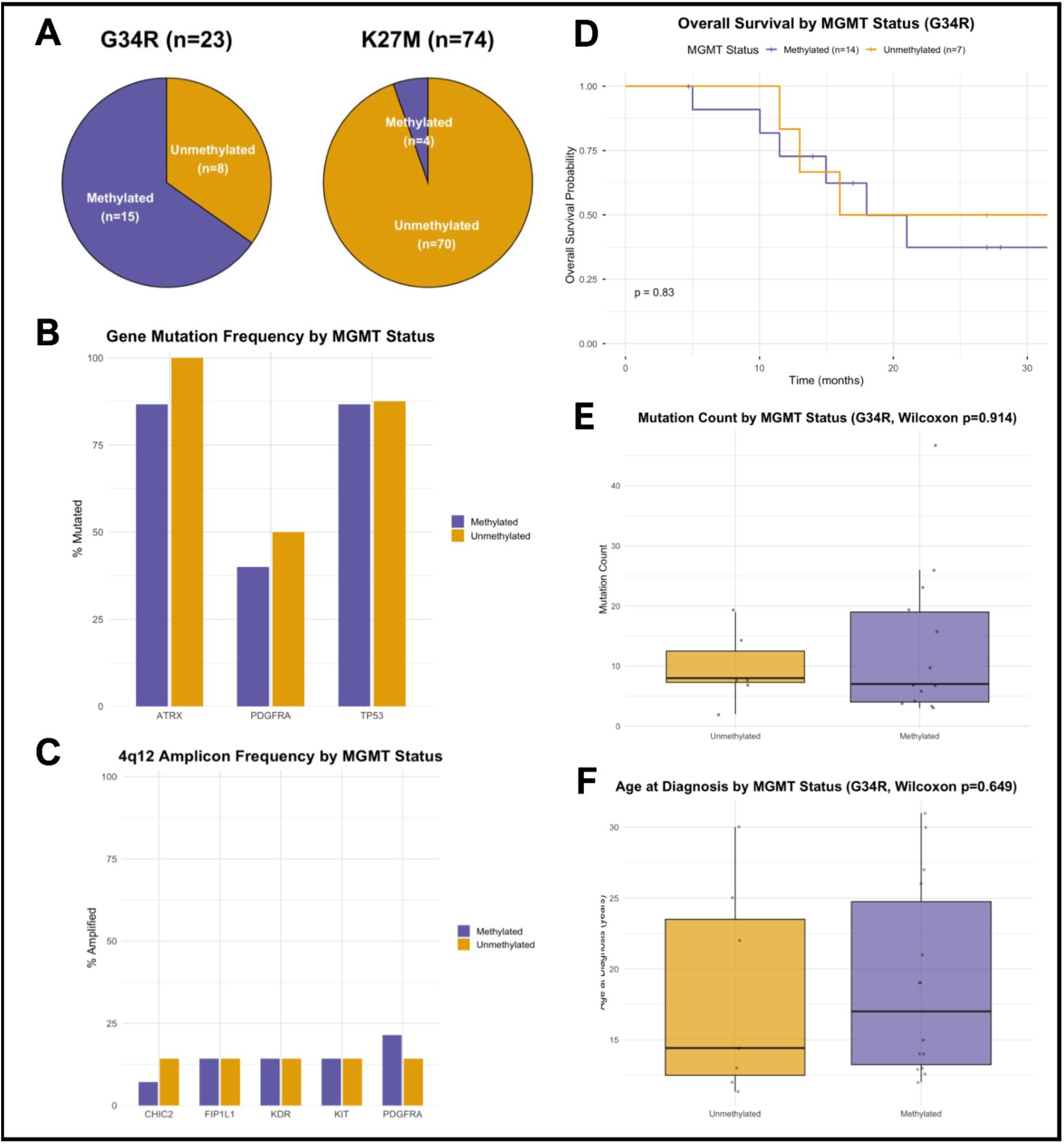
*MGMT* promoter methylation status and associated molecular and clinical features in *H3-K27M* and *H3-G34R* mutated gliomas. **(A)** Proportion of *MGMT*-methylated and -unmethylated samples in the *H3-G34R* (n=23) and *H3-K27M* (n=74) glioma cohorts. **(B)** Mutation frequency of *ATRX, PDGFRA*, and *TP53* by *MGMT* status within the *H3-G34R* cohort (methylated n=15, unmethylated n=8). **(C)** Copy-number amplification frequency at the chromosome 4q12 locus (*PDGFRA, KIT, CHIC2, FIP1L1, KDR*) by *MGMT* status within the *H3-G34R* glioma cohort. **(D)** Kaplan-Meier overall survival curves by *MGMT* status within the *H3-G34R* glioma cohort; log-rank test. One patient (GLSS-HK-0004) with discordant *MGMT* status across primary and recurrent samples was excluded from this comparison (methylated n=14, unmethylated n=7). **(E)** Mutation count by *MGMT* status within the *H3-G34R* glioma cohort; Wilcoxon rank-sum test. **(F)** Age at diagnosis by *MGMT* status within the *H3-G34R* glioma cohort (methylated n=14, unmethylated n=7, reflecting the same patient exclusion as **Panel D**); Wilcoxon rank-sum test. K27M is shown only as the overall methylated/unmethylated proportion (**Panel A**); the small number of *MGMT*-methylated K27M samples (n=4) was not meaningful enough for the group-level mutation, CNA, survival, and age comparisons shown for H3G34R (**Panels B–F**). See **Supplementary Table 1 and 2** for individual sample data.

Within the *H3-G34R* glioma cohort, mutation frequency of *ATRX, PDGFRA*, and *TP53* did not differ substantially between the *MGMT*-methylated (n=15) and *MGMT*-unmethylated (n=8) samples (*ATRX* 86.7% vs. 100%; *PDGFRA* 40.0% vs. 50.0%; *TP53* 86.7% vs. 87.5%; **Figure 4B**). Likewise, copy-number amplification frequency at the chromosome 4q12 locus (*PDGFRA, KIT, CHIC2, FIP1L1, KDR*) was similar between these subgroups, with a slightly higher *PDGFRA* amplification frequency among methylated samples (20.0% vs. 12.5%; **Figure 4C**).

Overall survival was not significantly different across the *MGMT*-methylated and unmethylated subgroups (log-rank test, p=0.83; **Figure 4D**). Mutation count (Wilcoxon rank-sum test, p=0.914; **Figure 4E**) and age at diagnosis (Wilcoxon rank-sum test, p=0.649; **Figure 4F**) also did not differ significantly across *MGMT* subgroups within the *H3-G34R* glioma cohort. One patient (GLSS-HK-0004) had both a primary and a recurrent *H3-G34R* glioma with unique genomic and epigenetics profiles as well as discordant *MGMT* status (astrocytoma, methylated; glioblastoma, unmethylated) despite a single, identical overall survival outcome. This patient was excluded from the survival comparison and age to avoid representing them multiple times, across both subgroups (methylated n=14, unmethylated n=7 for this comparison only). Overall, no significant molecular or clinical correlates of *MGMT* methylation status were identified within the *H3-G34R* glioma cohort in this exploratory analysis; the marked difference in methylation rate between cohorts (**Figure 4A**) remains the most notable finding.

### Cell-of-Origin Characterization among *H3-G34R* and *H3-K27M* gliomas

To evaluate the proposed cell-of-origin models for *H3-G34R* and *H3-K27M* glioma, we examined the mRNA expression levels for key markers for both of their lineage markers (**Figure 5A**). *SOX10* and *OLIG2*, are the primary markers of the oligodendrocyte precursor cell (OPC) lineage, which has been proposed as the cell of origin for *H3-K27M*-mutated DMG [12]. *GSX2, DLX1, DLX2,* and *FOXG1*, are markers of the ganglionic eminence (GE)/interneuron progenitor lineage proposed as the cell of origin for *H3-G34R*-mutated DHG [9].

**Figure 5.**
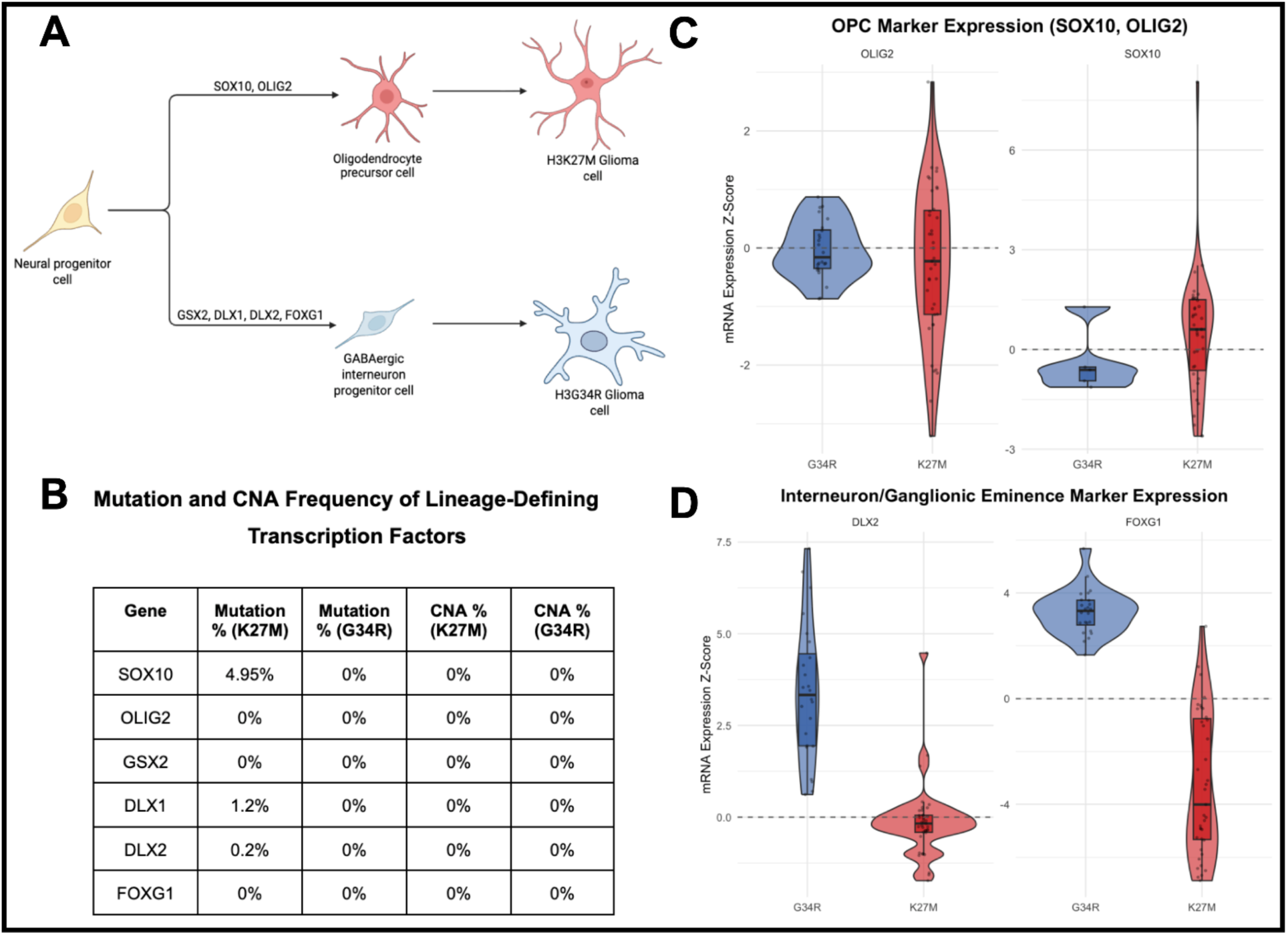
Cell-of-origin characterization of *H3-G34R* and *H3-K27M* mutant glioma. **(A)** Proposed developmental origin of each tumor type. *H3-K27M* glioma is proposed to arise from an oligodendrocyte precursor cell (OPC), marked by *SOX10* and *OLIG2* alterations while *H3-G34R* glioma is proposed to arise from a GABAergic interneuron/ganglionic eminence progenitor, marked by *GSX2, DLX1, DLX2,* and *FOXG1* alterations (Created with BioRender.com). **(B)** Mutation and copy-number alteration frequency of the six lineage-defining genes. **(C)** Expression of OPC lineage markers OLIG2 and SOX10 in *H3-G34R* and *H3-K27M* glioma samples, shown as mRNA expression z-score. G34R (RNA-seq, RSEM) and K27M (Agilent microarray) are shown on independent scales given differing expression profiling platforms; values are interpreted within, but not compared in magnitude across, cohorts. **(D)** Expression of GE/interneuron lineage markers DLX2 and FOXG1, shown and analyzed as in **Panel C**. GSX2 and DLX1 are not shown due to near-complete absence of *H3-K27M* glioma expression data for these genes (1 of 39 samples); see **Supplementary Table 3** for individual-sample data.

Mutation and copy-number status of these six genes was examined (**Figure 5B**). Except for *SOX10*, which was recurrently mutated in 4.95% of *H3-K27M* glioma samples as previously reported (**Figure 2A**), all six genes were mutated or CNA-altered in less than 2% of samples in either cohort, or several showed no alterations at all. This near absence of genetic alteration is consistent with these genes’ proposed role as epigenetically inherited markers of developmental lineage rather than somatically acquired drivers [58].

Expression of the OPC markers *SOX10* and *OLIG2* was compared within each cohort against a reference z-score of 0 (Figure 65). *SOX10* expression was moderately elevated in *H3-K27M* gliomas (median z-score = 0.488, n=39) while also being reduced below baseline in *H3-G34R* gliomas (median z-score = -0.384, n=5), indicating active suppression of this OPC marker outside its proposed matching lineage rather than simple absence of elevation. However, given the small number of G34R samples with reported *SOX10* expression data (n=5), this result should be interpreted with caution. *OLIG2* expression was not meaningfully different from baseline in either glioma cohort (K27M: median z-score = -0.262, n=39; G34R: median z-score = -0.029, n=24). Expression of the GE/interneuron markers *DLX2* and *FOXG1* showed the converse pattern within their respective glioma cohorts (**Figure 5D**). Both genes were highly elevated in *H3-G34R* (DLX2: median z-score = 3.416, n=24; FOXG1: median z-score = 3.28, n=24). The latter also showed markedly reduced expression in *H3-K27M* gliomas (median z-score = -3.25, n=39), again indicating active suppression rather than mere absence of elevation. *DLX2* expression did not differ substantially from the baseline in *H3-K27M* gliomas (median z-score = -0.127, n=39). Other prominent ganglionic eminence (GE)/interneuron progenitor markers such as *DLX1* and *GSX2* showed notable upregulation in *H3-G34R* gliomas (*DLX1*: median z-score = 3.420, n = 24; *GSX2*: median z-score = 5.142, n = 24; **Supplementary Table 3**). *GSX2* and *DLX1* expression could not be evaluated in *H3-K27M* gliomas due to near-complete absence of data for these two genes in the available Agilent microarray cohort (1 of 39 samples with a reported value for each gene); individual case-level data for this sample are provided in **Supplementary Table 3**.

Together, these findings support a model in which *H3-G34R* and *H3-K27M* gliomas arise from distinct, non-overlapping progenitor lineages.

## Discussion

This study is the first study in the existing literature that directly compares the mutational, epigenetic, and clinical correlates of *H3-G34R* and *H3-K27M* gliomas. Although both are associated with genomic and epigenetic dysregulation, their respective mechanisms are quite different. Our study provided a way to investigate which features of these tumors are specific to the disease, as compared to traits that are shared hallmarks of oncohistone-driven gliomagenesis more broadly.

Our demographical and anatomic analysis reproduced the expected clinical findings for both *H3-G34R* and *H3-K27M* gliomas. Anatomically, the former glioma is characterized by dominant hemispheric presence, with the latter more associated with midline presence, with meaningful non-pontine representation as well [59]. Our analysis revealed similar findings, showing that *H3-K27M* gliomas are more likely to expand outside of their characteristic regions than their *H3-G34R* counterparts, being consistent with the idea that they might be more aggressive. Both findings agree with established clinical patterns, and this supports the idea that our pooled cBioPortal/pedcBioPortal cohort is a reasonably representative sampling frame for the genomic comparisons that followed.

Characterization of the *H3-G34R* glioma genomic landscape established a unique mutational profile, consistent with previous findings, identifying *TP53* (94.1%), *ATRX* (81.4%), and *PDGFRA* (54.9%) as top-mutated genes [9,19]. Alongside this copy-number and structural variant data revealed comparable findings to *H3-K27M* glioma. Characterization of the *H3-K27M* mutational landscape identified a recurrently altered gene panel consisting of *TP53* (62.2%), *ATRX* (21.2%), *PPM1D* (19.3%), *PIK3CA* (14.9%), *NF1* (13.3%), and *PDGFRA* (10.8%). Interestingly enough, our observed *ACVR1* mutation frequency (∼4%) was substantially lower than rates reported in earlier *H3-K27M* DMG-focused literature (∼20–25%) [12]. We attribute this discrepancy to cohort composition – namely, our cohort consisted of gliomas harboring *H3-K27M* mutation across different midline and even cerebral regions, yet the *ACVR1* mutation is specifically associated with the *H3.1-K27M* mutation, which is known to be concentrated in pons-related cases, specifically DIPG [60]. These findings illustrate the vast anatomic and genetic heterogeneity existing in both tumors.

The direct comparison between cohorts confirmed that *H3-G34R* and *H3-K27M* have distinct clinical and molecular characteristics. Patients with *H3-G34R* gliomas were diagnosed at a significantly older age than those with *H3-K27M* gliomas (Wilcoxon p=5.7×10^-12^), consistent with previous reports that hemispheric and midline gliomas preferentially manifest in young adults and children, respectively [61]. The two cohorts differed significantly in overall survival (log-rank p=0.00015), with *H3-G34R* glioma patients significantly more likely to live longer than their *H3-K27M* glioma patient counterparts, indicating that the latter is more aggressive. This interpretation of *H3-K27M* gliomas being more aggressive is consistent with our previous findings about anatomic location, and the disparity in overall survival matches existing data in other studies [3,19].

Recurrent gene mutation comparisons reinforced largely non-overlapping profiles between the two cohorts. These differences were made more explicit through each cohort’s respective GO biological processes analysis, with *H3-G34R*-associated mutations converging on processes such as cell cycle regulation, cellular senescence, and epigenetic regulation of gene expression, and *H3-K27M*-associated mutations converging on PI3K/Akt signal transduction, oligodendrocyte differentiation, and negative regulation of neurogenesis. Copy-number analysis identified a shared chromosome 4q12 amplicon consisting of *PDGFRA, KIT, CHIC2, FIP1L1*, and *KDR*, with amplification rates at modestly higher frequency in *H3-G34R* gliomas across all five genes. Structural variant data between both cohorts was much more similar than the mutational data. An important caveat here is that structural variant frequencies in *H3-G34R* gliomas (∼6–7%) were low enough to plausibly reflect background noise rather than true recurrent partners, which resulted in our more cautious interpretation of those data.

*MGMT* promoter methylation status was assessed in both cohorts, with a substantially larger methylated fraction identified in *H3-G34R* than in *H3-K27M* gliomas (15/8 vs 4/70 methylated/unmethylated; Fisher’s Exact Test: p=8.3×10^-9^). The *H3-K27M* glioma cohort had too few *MGMT+* samples and was underpowered for meaningful group-level statistical comparison, so instead was reported at the level of individual cases (**Supplementary Table 1-2**). No strong or consistent signal distinguishing methylated from unmethylated tumors emerged in either cohort within the constraints of this sample size. This analysis should be read as hypothesis-generating rather than conclusive, particularly for *H3-K27M* gliomas.

Finally, cell-of-origin characterization investigated mRNA expression levels across a panel of lineage-defining transcription factors for *H3-G34R* and *H3-K27M* gliomas (*GSX2, DLX1, DLX2, FOXG1*, and *SOX10, OLIG2*, respectively) in both cohorts. Our results are consistent with established cell-of-origin models established for these tumors indicating that these genes function as epigenetically inherited markers of developmental identity rather than somatically targeted drivers [9,12,13,19]. Expression analysis was interpreted within each cohort, due to the separate sequencing methods; namely, the RNA-seq for *H3-G34R* gliomas and Agilent microarray platform for *H3-K27M* gliomas. Our results found *FOXG1* and *DLX2* to be elevated in *H3-G34R* and suppressed in *H3-K27M*, while *SOX10* was elevated in *H3-K27M* and reduced in *H3-G34R*, with the *SOX10, H3-G34R* result drawn from a small sample (n=5) and interpreted cautiously. We also found *DLX1* and *GSX2* to be meaningfully upregulated in *H3-G34R* glioma tumors but had too few *H3-K27M* glioma samples (n=1) to meaningfully interpret any patterns for those genes in those tumors. Together, these results support a model in which *H3-G34R* and *H3-K27M* gliomas arise from and retain the transcriptional signature of distinct progenitor populations – an OPC-like lineage in the case of *H3-K27M* and a GE/interneuron-like lineage for *H3-G34R* – rather than converging on a shared cell of origin.

The prominent co-occurrence of *TP53* and *ATRX* mutation in both *H3-K27M* and *H3-G34R* glioma cohorts is consistent with a well-established signature of genomic instability and alternative lengthening of telomeres (ALT) in high-grade glioma [62,63], and our finding here does not necessarily extend what is already known about this relationship in either *H3-G34R* or *H3-K27M* gliomas specifically.

Another indication of genomic and cell cycle-related instability is the reported sensitivities of these high-grade gliomas to CDK4/6 therapy, especially *H3-G34R* [19,53,64]. Interestingly, we observed CDK6 amplification in only 2.6% of *H3-G34R* glioma samples (**Figure 1C**), which is considerably lower than the rates reported in prior studies [65]. Additionally, *CDKN2A/CDKN2B* homozygous deletion is a prominent feature in *H3-G34R* gliomas, but our analysis found this to be the case in only ∼5% of such tumors (**Figure 1C**) [65,66]. However, studies have generally found that CDK4/6 inhibitor sensitivity does not track cleanly with *CDK4/6* copy-number status across glioma types [66–68]. This means that the low *CDK6* amplification frequency in our *H3-G34R* glioma cohort does not by itself argue against CDK4/6 inhibition as a rational strategy. Given that "G1/S phase transition" emerged independently as one of the most highly enriched GO terms among recurrently altered genes in the *H3-G34R* glioma cohort (**Figure 1D**), it is plausible that CNAs in other genes involved in the G1/S checkpoint underlie the CDK4/6 inhibitor sensitivity reported elsewhere.

A related, *H3-K27M* glioma-specific vulnerability is suggested by recurrent mutations within the PI3K/AKT signaling pathway. GO enrichment analysis identified "PI3K/Akt signal transduction" as the single most prominent biological process associated with *H3-K27M*-mutant alterations (**Figure 2D**), and this prominence is reflected when examining recurrently mutated genes such as: *PIK3CA* (14.9%), *NF1*(13.3%), *PDGFRA* (10.8%), *PIK3R1* (7.8%), *PTEN* (5.7%), and *FGFR1* (4.5%) (**Figure 2B**). *PTEN* is the primary molecular brake on the PI3K signaling pathway [69], *NF1* is involved in this pathway through its role upstream in RAS/PI3K signaling [70], and *PDGFRA* encodes an RTK whose activation directly feeds PI3K/AKT [71,72]. Together, these mutations illustrate how multiple independent alterations along a single signaling cascade, can lead to oncogenic growth, and suggests PI3K/AKT/mTOR inhibition as a rational, therapeutic strategy specifically in *H3-K27M* glioma, one already under early preclinical and clinical investigation in diffuse midline glioma [73].

Beyond pathway-specific inhibition, the most frequent focal amplification identified in both *H3-G34R* and *H3-K27M* glioma cohorts involved the chromosome 4q12 amplicon, which includes *PDGFRA, KIT*, and *KDR* – three receptor tyrosine kinases (RTKs) co-amplified on the same locus – along with two nearby genes (*CHIC2, FIP1L1*) that are associated with RTKs (**Figure 3C**). These recurrent alteration patterns have led to RTK-targeted therapy becoming a strongly supported preclinical rationale for the treatment of these tumors [74–76]. Although this rationale makes sense intuitively, single-agent RTK inhibitors have historically shown limited clinical efficacy in human trials due to extensive tumor heterogeneity, compensatory signaling loops, and the blood-brain barrier [76]. Presenting this rationale alongside our own amplicon frequency data, generated from a pooled contemporary cohort, adds concrete support to the case for investigating either multi-model RTK inhibition or RTK inhibition in conjunction with other therapeutic options. Taken together with the *TP53/ATRX* alterations and G1/S findings above, our data are broadly consistent with four existing therapeutic rationales in this disease: PARP and Chk1/2 inhibition motivated by *TP53/ATRX*-driven genomic instability [19,77,78], CDK4/6 inhibition suggested by G1/S pathway dysregulation [19,53,64], PI3K/AKT inhibition prompted by convergent pathway mutation in H3-K27M gliomas [73], and RTK inhibition suggested by chromosome 4q12 amplification [74–76].

Our lab has reported a preclinical effect of the imipridone drug class that includes ONC201 and ONC206 towards reduction of MGMT as well and EZH1 and EZH2 protein expression in *H3-K27M* gliomas [79–82]. This dual effect is particularly relevant to two central findings of this analysis. First, *MGMT* promoter methylation is extremely rare in *H3-K27M* gliomas relative to *H3-G34R* gliomas (4/74, 5.4% vs 15/23, 65.2%; **Figure 4A**). *MGMT*-unmethylation is generally associated with poorer response to temozolomide, the primary alkylating agent used as standard-of-care for the treatment of glioblastoma tumors, and this finding indicates that most *H3-K27M* glioma tumors are intrinsically resistant to this therapy [83,84]. Second, EZH2 reduction directly targets the defining molecular lesion of *H3-K27M* glioma itself. Mechanistically, it is the mutant methionine’s high-affinity sequestration of PRC2 via binding the EZH2 domain that results in the genome-wide loss of H3K27me3 (**Figure 2**). However, recent studies have found that ONC201 can increase H3K27me3 independent of EZH2 [85], a finding that is considered generally favorable for the treatment of *H3-K27M* gliomas [86]. In this sense, ONC201/ONC206 represent a mechanistically distinct addition to the rationales outlined above: where PARP/Chk1-2, CDK4/6, PI3K, and RTK inhibition are motivated primarily by co-occurring genomic alterations, imipridone-class agents offer a strategy aimed at the epigenetic mechanism that defines *H3-K27M* glioma.

The main limitations in this study lie in the limited sample sizes for much of the analysis. This study is also limited by its retrospective, cross-sectional design, which gathers data from publicly deposited glioma cohorts profiled on different sequencing platforms and gene panels. As such, much of the clinical annotation, particularly *MGMT* methylation status, was available for only a minority of samples in both cohorts, and other measures, such as differentiation marker expression, were not even comparable due to the different methods of measurement. Additionally, the therapeutic rationales proposed here are drawn entirely from genomic and transcriptomic inference; none have been functionally validated in cell or animal models as part of this study and should be treated as hypothesis-generating rather than established. In the future, it will be important to investigate large molecular profiling databases with extensive repositories on the genetic, epigenetic, and clinical attributes of *H3-G34R* and *H3-K27M* glioma patients. This could provide a more detailed narrative surrounding the biological drivers of *H3-G34R* and *H3-K27M* gliomagenesis by investigating chromatin remodeling, modified histone analysis, and global *MGMT* methylation patterns by utilizing transcriptomic and sequencing analyses. Access to such data could lead to a more robust understanding of the underlying biology behind these aggressive gliomas with epigenetic mutations and other alterations to further elucidate the mechanisms through which these underlying biological factors induce gliomagenesis.

Taken together, our study suggests that, despite their shared status as oncohistone-driven, poor-prognosis high-grade gliomas, *H3-G34R* and *H3-K27M* gliomas are best understood as distinct disease entities arising from distinct progenitor lineages, with distinct – albeit partially overlapping – pathogenetic mechanisms and therapeutic vulnerabilities. Specific precision oncology therapeutic vulnerabilities across *H3-G34R* and *H3-K27M* glioma subtypes, based on their mutational and copy-number profile is suggested as well as differentiation-related growth pathways that require further study as potential loci for therapy.

## Acknowledgements

W.S.E-D. is an American Cancer Society Research Professor and is supported by the Mencoff Family University Professorship at Brown University.

---

**Supplementary Table 1:**
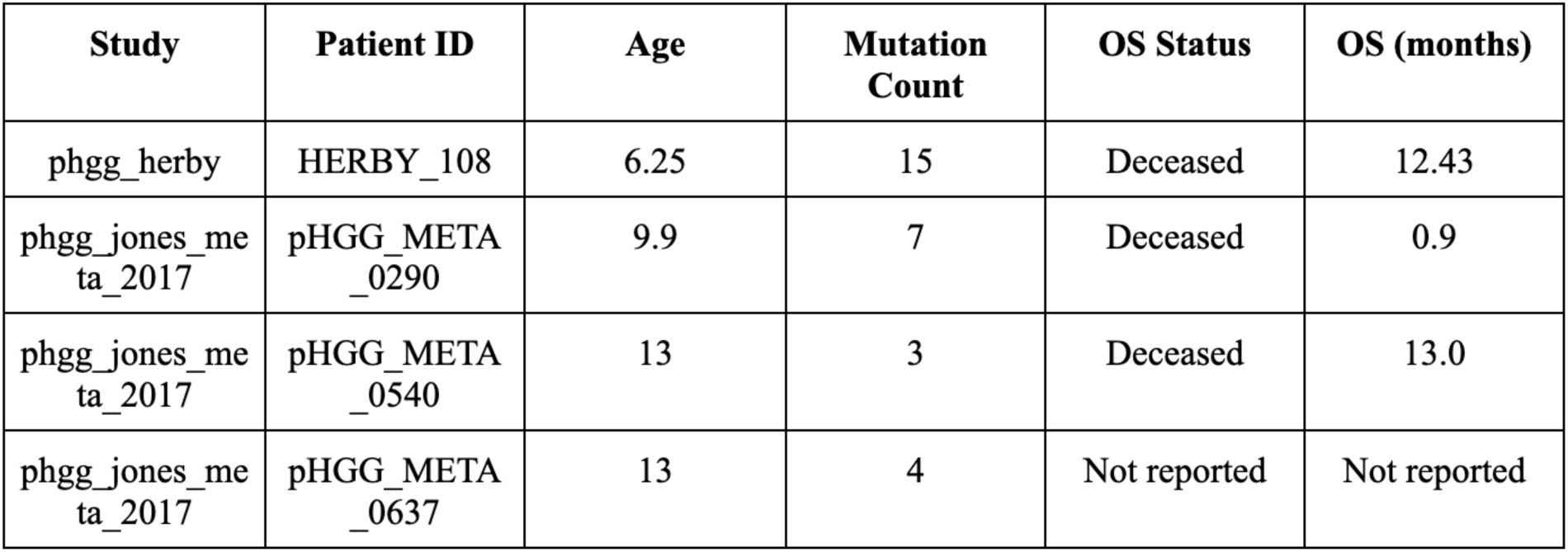
Age, Mutation Count, OS in the MGMT+ K27M subgroup.

| **Study** | **Patient ID** | **Age** | **Mutation Count** | **OS Status** | **OS (months)** |
| --- | --- | --- | --- | --- | --- |
| phgg_herby | HERBY_108 | 6.25 | 15 | Deceased | 12.43 |
| phgg_jones_meta_2017 | pHGG_META_0290 | 9.9 | 7 | Deceased | 0.9 |
| phgg_jones_meta_2017 | pHGG_META_0540 | 13 | 3 | Deceased | 13.0 |
| phgg_jones_meta_2017 | pHGG_META_0637 | 13 | 4 | Not reported | Not reported |

**Supplementary Table 2 :**
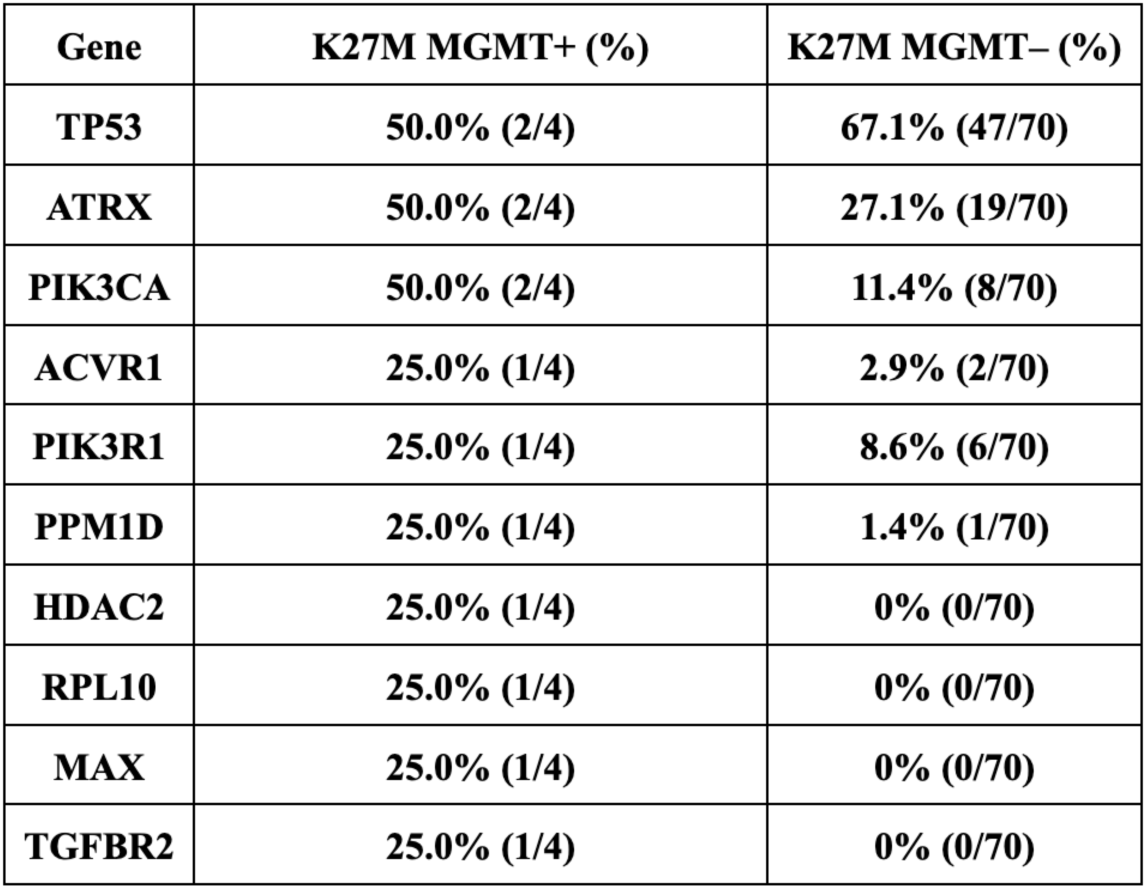
Recurrent mutations by K27M MGMT cohort.

| **Gene** | **K27M MGMT+ (%)** | **K27M MGMT– (%)** |
| --- | --- | --- |
| **TP53** | **50.0% (2/4)** | **67.1% (47/70)** |
| **ATRX** | **50.0% (2/4)** | **27.1% (19/70)** |
| **PIK3CA** | **50.0% (2/4)** | **11.4% (8/70)** |
| **ACVR1** | **25.0% (1/4)** | **2.9% (2/70)** |
| **PIK3R1** | **25.0% (1/4)** | **8.6% (6/70)** |
| **PPM1D** | **25.0% (1/4)** | **1.4% (1/70)** |
| **HDAC2** | **25.0% (1/4)** | **0% (0/70)** |
| **RPL10** | **25.0% (1/4)** | **0% (0/70)** |
| **MAX** | **25.0% (1/4)** | **0% (0/70)** |
| **TGFBR2** | **25.0% (1/4)** | **0% (0/70)** |

**Supplementary Table 3 :**
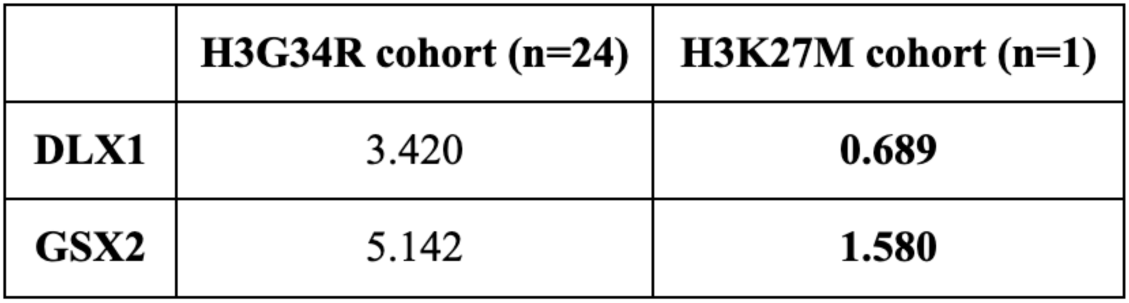
DLX1 and GSX2 mRNA expression.

